# Regulations of mitochondrial DNA repair by poly(ADP-Ribose) Polymerase 1

**DOI:** 10.1101/2020.07.13.200055

**Authors:** Geoffrey K. Herrmann, Y. Whitney Yin

## Abstract

Formation of a repair enzyme complex is beneficial to DNA repair. Despite the fact that mitochondrial base excision repair (mtBER) enzymes DNA polymerase gamma (Pol γ) and poly(ADP-ribose) polymerase 1 (PARP1) were found in the same complex, the functional role of the interaction in mtBER has not been characterized. We report studies that PARP1 regulates Pol γ activity during DNA repair in a metabolic cofactor NAD^+^ (nicotinamide adenosine dinucleotide)-dependent manner. In the absence of NAD^+^, PARP1 completely inhibits Pol γ, while increasing NAD^+^ level to physiological concentration enables Pol γ to resume maximum repair activity. Pol γ is PARylated when bound to DNA repair intermediates, and PARylation is essential for Pol γ repair activity. The PARP1 inhibitor Olaparib that abolishes PARP1 catalytic activity suppresses Pol γ gap-filling synthesis at physiological concentrations of NAD^+^, suggesting inhibiting PARP1 activity would increase mtDNA mutations. Because NAD^+^ cellular levels are linked to metabolism and to ATP production via oxidative phosphorylation, our results suggest that mtDNA damage repair is correlated with cellular metabolic state and integrity of the respiratory chain. Our results revealed a molecular basis of drug toxicity from prolonged usage of PARP1 inhibitors in treating cardiac dysfunctions

## INTRODUCTION

The oxidative environment of mitochondria results in more damage on mitochondrial DNA (mtDNA) than its nuclear counterpart (*1, 2*). Most of mtDNA oxidative damage is restored via base excision repair (mtBER), which, like nuclear BER, can be functionally grouped into three distinct steps: 1) Lesion recognition, 2) Gap tailoring, and 3) DNA synthesis/ligation (*3*). Although mtBER Lesion recognition is similarly to nuclear BER, the pathway subsequently diverges. In contrast to specialized DNA polymerases in the nucleus, mitochondrial DNA polymerase (Pol γ) plays dual roles in DNA replication and repair (*4*), but regulation of Pol γ repair vs. replication has not been illustrated. Pol γ does not appear suitable for canonical nuclear BER because of its low-efficiency synthesis on a single-nucleotide gapped DNA and lack of strand displacement DNA synthesis ability (*5*), both of which are essential for single-nucleotide (SN) and long-patch (LP) BER, respectively. However, Pol γ gap-filling synthesis is enhanced by the mitochondrial specific endo/exonuclease G (EXOG), which precisely excises two-nucleotides from the 5’-end of the gap of duplex DNA, thereby creating an optimal substrate for Pol γ (*5-7*). This mtBER product resembles those from nuclear LP-BER but is produced by a different mechanism.

Poly(ADP-Ribose) (PAR) polymerase 1 (PARP1) is a first responder to DNA strand breaks and serves as a key scaffolding protein in nuclear BER (*8, 9*). By catalyzing poly(ADP-ribosylation) on itself (auto-PARylation) and targeting proteins (trans-PARylation) using substrate NAD^+^, PARP1 serves as a scaffold in recruiting other repair enzymes to the lesion site (*10, 11*). Despite a definitive role of PARP1 in nuclear BER, both positive and negative roles of PARP1 in mtDNA repair have been reported. Knockdown of PARP1 gene reduces DNA damage (*12*) but a PARP1 deletion increasing DNA damage under oxidative conditions (*13*), and the PARP1 inhibitor augments UV-induced damage on mtDNA (*14*).

Despite its functional ambiguity in mtBER, PARP1 was found in complex with other mtDNA repair enzymes apurinic/apyrimidinic enduonuclease 1 (APE1), Pol γ, EXOG, and Ligase III in mitochondrial extract (*15*). Futher, Pol γ was found to interact with PARP1 directly and be PARylated in cardiomyocytes during parasite infection (*12*). Formation of a repair enzyme complex, or repairosome, not only increases repair efficiency but also prevents premature release of DNA repair intermediates that could be more detrimental to the cells than the original lesion (*16*). Functional and structural details of PARP1-Pol γ interaction and its impact on mtDNA repair are unknown.

Studies of PARP1’s activities in mitochondria were conducted in cells or animals, where DNA repair is complicated by PARP1 activation-induced NAD^+^ reduction or depletion, particularly when PARP1 was pathologically overexpressed PARP1 in cardiomyocytes (*17*). NAD^+^ is a major cofactor for glucose and fatty acids metabolism; the resulting NADH is converted back to NAD^+^ by the oxidative phosphorylation (OXPHOS) electron transfer chain coupled with ATP synthesis. Depletion of NAD^+^ can negatively impact metabolism and ATP production.

To begin dissecting PARP1 function in mtDNA repair from its role in metabolism, we conducted biochemical studies to investigate PARP1 function on Pol γ functions. We report here that PARP1 differentially affects Pol γ in replication and repair. While having little impact on Pol γ during replication, PARP1 regulates Pol γ repair activity in a NAD-dependent manner. Pol γ is a substrate for PARP1, and PARylation is essential for PARP1 regulation of Pol γ repair activities. Our studies thus revealed a crosstalk between mtDNA repair and cellular metabolism.

## RESULTS

### PARP1 interacts with Pol γ on DNA repair intermediates

To recapitulate the interaction of PARP1 and Pol γ as had been captured by co-immunoprecipitation (co-IP) (*12*), we used purified PARP1 and Pol γ with or without a 75-nt dumbbell gapped DNA that mimics a mtDNA BER intermediate (Fig. EV1A). Pol γ is comprised of a catalytic subunit Pol γA that houses a polymerase (*pol*) and an exonuclease (*exo*) active sites and a dimeric accessory subunit Pol γB. Pol γ holoenzyme was formed by mixing Pol γA and Pol γB and purified as a complex of Pol γAB_2_.

Binary complexes of Pol γ -DNA, PARP1-DNA, and PARP1-Pol γ were first measured using nano-isothermocalorimetry (nano-ITC). At 25°C, the enthalpy for PARP1-DNA binary complex was -58.4 kcal/mol, and the dissociation constant (K_d_) was calculated to be 36.4 nM. There was no measurable ΔH for Polγ binding to DNA at 25°C (Fig. EV1B), which simplified evaluation of the Pol γ-PARP1-DNA ternary complex. The calculated K_d_ for the ternary complex was 111.1 nM (Fig. EV1C) with enthalpy ΔH_PARP1:Pol γ:DNA_ was -61.4 kcal/mol similar to PARP1-DNA binary complex. This results suggest formation of PARP1-Pol γ -DNA ternary complex, and binding is entropically driven (Fig. EV1D). The lower affinity of the ternary complex, in which PARP1 and Pol γ are simultaneously bound to the DNA, relative to either binary complex, maybe due to steric hindrance at the binding of Pol γA *pol* and PARP1 Zn2 domain to, respectively, to the 3’- and the 5’-end of the DNA gap (*18-20*).

As multi-component binding constants are difficult to deconvolute from ITC data, we performed electrophoretic mobility shift assays (EMSAs) to visualize complex formation directly. EMSA supershift assays were performed by titrating increasing amounts of PARP1 into a constant quantity of Pol γ-^32^P-DNA complex. A super-shifted species emerged that migrated differently from either PARP1-DNA or Pol γ-DNA complexes. The quantity of this species is positively correlated with PARP1 concentration (Fig. EV2A, EV2B). To identify the components in the supershifted species, Western blot was performed using antibody against Pol γ? When PARP1 is added, the supershifted band contains Pol γ. Because the reaction was under substoichiometry of Pol γ to DNA (2:3 molar ratio) thus only one Pol γ is bound to each DNA substrate, therefore, the supershifted Polγ-DNA complex most likely also contains PARP1. The supershifted species dissociates upon addition of NAD back to Pol γ-DNA binary complex (Fig. EV2C, lane 4-5), and addition of Olaparib prevents the dissociation (Fig. EV2C, lane 10), suggesting the supershifted species is a Pol γ-PARP1-DNA complex and its stability is reduced by PARylation.

### Pol γ trans-PARylatin on DNA repair intermediate

To examine whether Pol γ is a substrate for PARP1, PARP1 PARylation of Pol γ was analyzed by Western Blot. As PARP1 PARylation is stimulated >1000-fold in presence of DNA (*21*), we utilized two DNA constructs: the dumbbell 3-nt gapped DNA substrate used for ITC and EMSA analyses that is sufficient to accommodate both PARP1 and Pol γ, and an 8bp self-annealing DNA duplex that can stimulate PARP1 activity but only long enough to bind PARP1 alone (*22*). These substrates allow evaluation of Pol γ PARylation free in solution (8bp duplex DNA) and in close vicinity to PARP1 (3-nt gapped DNA).

PARylation reaction products could potentially contain both PARylated as well as unmodified PARP1 and Pol γ. As heterogeneous PAR chains yield smear on a denatured PAGE gel where the PARylated Pol γ and PARP1 overlap, Western blot was used to distinguish each PARylated species using antibodies against PARP1 and Pol γ. For each reaction with a specific DNA substrate, anti-PARP and anti-PARP1 antibodies were sequentially employed (Methods). Auto-PARylation of PARP1 was equally robust on both DNA substrates, which was clearly evident at 20 µM NAD^+^ (most clearly by the loss of the discrete band for the unmodified species). Auto-PARylation increased with increasing concentrations of NAD^+^, plateauing around a physiological concentration of NAD^+^ at 200 µM (Fig. 1B, 1D) (*23, 24*). By contrast, trans-PARylation of Pol γ differed significantly on the two DNA substrates. On the 8bp duplex, only a small amounts of short PAR chains were observed and only at 2000 µM NAD^+^ (Fig. 1A), whereas on the gapped DNA, Pol γ PARylation began at 20 µM NAD^+^, increasing sharply at 200 µM NAD^+^ where ∼50% of Pol γ were modified (Fig. 1C). These results suggest that at physiological concentrations of NAD^+^, Pol γ was PARylated more readily when bound to the same gapped DNA as PARP1 than when free in solution. Interestingly, auto-PARylated PARP1 did not increase on 8bp DNA when Pol γ was not PARylated, suggesting that PARP1 catalyzes auto-PARylation and trans-PARylation as two separate reactions and without preference.

**Figure 1.**
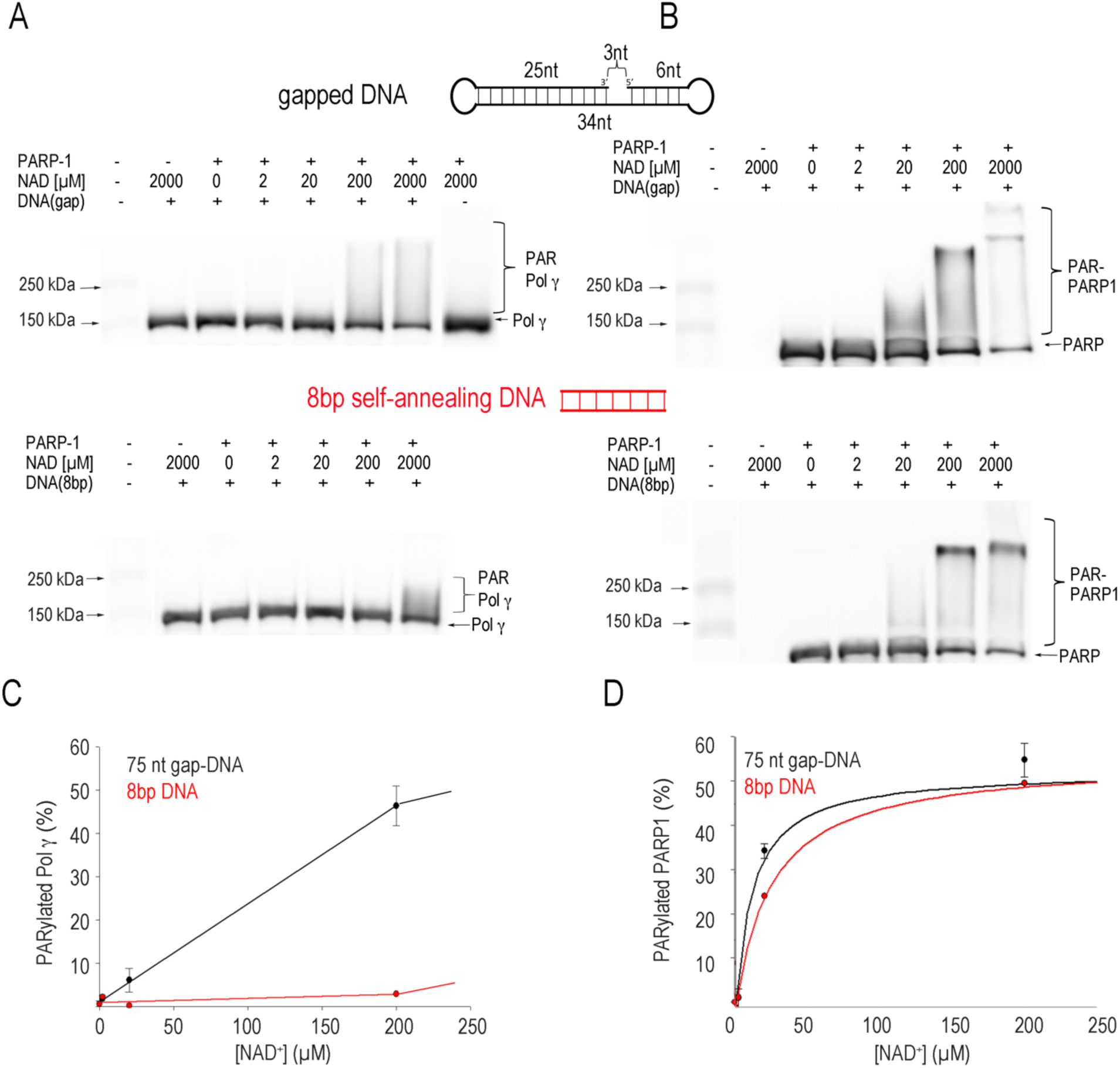
DNA dependent Pol γ trans-PARylation. A) Pol γ trans-PARylation on 75nt gapped- and 8bp DNA, respectively, and C) quantification. B) PARP1 auto-PARylation on 75 nt gapped- and 8bp DNA, respectively, and their quantification.

To evaluate whether PARP1 modifies any repair protein that displays affinity to gapped DNA, we tested human EXOG PARylation on the same gapped DNA and found no PARylation of EXOG (Supplementary Information and Fig. EV3). Taken together, the results indicate that Pol γ is a bona fide substrate for PARP1.

### PARP1 modulates Pol γ gap-filling activity in a NAD^**+**^**-dependent manner**

To examine the biological effects of PARP1 interaction with Pol γ and PARylation, we first examined Pol γ activities in DNA repair on a gapped DNA substrate that mimics the DNA intermediate following hEXOG excision (Fig. 2A) (*6, 7*).

**Figure 2.**
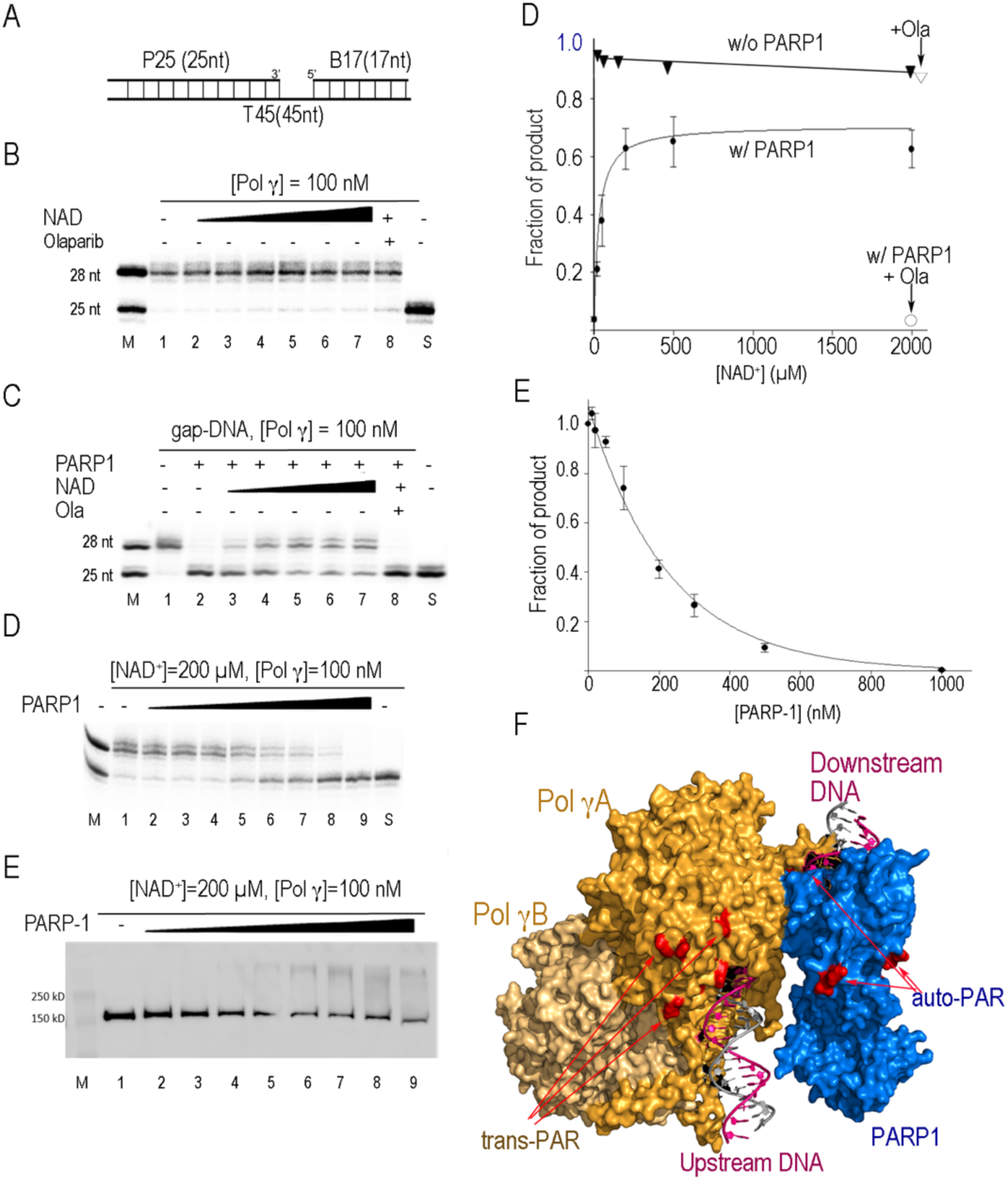
PARP1 regulates Pol γ gap-filling synthesis A) Substrate used in assays, B) Pol γ gap-filling synthesis in the absence (Lane 1) or presence of NAD^+^ (20 µM, 50 µM, 200 µM, 500 µM and 2000 µM, Lanes 2-7), and in the presence of Olaparib (10 µM, Lane 8). Lane M and S denote marker and substrate only lane, C) Same as in A, except 200 nM PARP1 is present. D) Pol γ (100 nM) gap-filling synthesis in presence of constant NAD^+^ (200 µM) and increasing concentration of PARP1 (10 nM, 20 nM, 50 nM, 100 nM, 200 nM, 300 nM, 500 nM, 1000 nM, Lanes 2-9). E) Western blot using antibody against Pol γ showing Pol γ PARylation and F) Quantification of Fraction of product formed in A), G) Quantification of D, H) PARylation sites identified by LC-MS/MS mapped on a model of PARP1 Pol γ-gap DNA complex composited from crystal structures. PARP1 (blue) is located at the 5’-end of the gap and Pol γ at the 3’-end where Pol γA (Orange) is in close contact with PARP1 and Pol γB (Light orange) is distal to the DNA gap. The PARylation sites detected are colored red.

Pol γ alone efficiently synthesized into the 3-nt DNA gap (Fig. 2B). However, addition of stochiometric amount of PARP1 completely inhibited Pol γ gap-filling activity (Fig. 2C, Lane 2). The diminished Pol γ activity is most likely due to PARP1 obstructing Pol γ from accessing the template in the DNA gap. To examine the effects of PARylation, NAD^+^ were titrated into the reaction mixture. As NAD^+^ concentration increases, beginning at 50 µM NAD^+^, Pol γ gap-filling activity was gradually recovered (Fig. 2C, Lane 3-7), and reached a maximum at 200 µM. Further increasing NAD^+^ to 2000 µM offers negligible benefit (Fig. 2F). NAD^+^ alone does not affect Pol γ activity (Fig. 2B, 2F), thus NAD^+^ must regulate PARP1/Pol γ interaction through PARylation.

Recent studies suggest that the human nuclear BER polymerase, Pol β, could be localized in mitochondria and therefore could participate in mtBER. We thus tested the effect of PARP1 on Pol β gap-filling synthesis. Similarly to results from Pol γ, Pol β activity is also inhibited by PARP1, and restored by the addition of NAD^+^ (Fig. EV4), consistent with the previous report (*25*). Regardless of which polymerase performs BER activity in mitochondria, its gap-filling synthesis is regulated by PARP1 and NAD^+^.

To confirm PARylation is a key regulator for PARP1-Pol γ interaction, we added to the reaction a PARP inhibitor, Olaparib, which competes with NAD^+^ thus obliterates PARP1 PARylation activity. Olaparib completely abolishes the NAD^+^ rescuing effect (Fig. 2C, Lane 8), confirming the catalytic activity of PARP1 is involved in regulation of Pol γ activity. These results show that PARP1 protests broken DNA and facilitates Pol γ gap-filling synthesis in the presence of NAD^+^, indicating PARP1 contribute positively to gap-filling activity in mtDNA BER.

### Excessive PARP1 halts Pol γ DNA repair activity

To address the reported PARP1’s negative impact, we examined Pol γ gap-filling activity with elevated concentration of PARP1 at constant physiological NAD^+^ level to mimic PARP1 overexpression under pathological conditions. Pol γ activity were examined at various PARP1:Pol γ molar ratios from 1:10 to 10:1 (Fig. 2D). Gap-filling product began to decrease at the molar ratio of 1:1 and continues exponentially with increasing PARP1 stoichiometry (Fig. 2G). Pol γ activity is completely halted at 10-fold molar excess of PARP1 to Pol γ (Fig. 2D, lane 9).

The excess molar ratio of PARP1 to Pol γ provides an opportunity to examine the order of auto-PARylation vs. trans-PARylation, i.e., whether PARP1 preferentially catalyzes auto-PARylation of itself prior to trans-PARylation of substrates with limited NAD^+^. Previous studies show that each PAR chain can contain about 200 ADP-ribose moieties (*26*) and each PARP1 molecule has at least 37 PARylation sites (*27*). If PARP1 preferentially catalyzes auto-PARylation, assuming 50% of PARP1 are active (Fig. 1B), at PARP1 concentration greater than 100 nM, auto-PARylation could consume all 200 µM NAD^+^ in the reaction thus leave no NAD^+^ for trans-PARylation of Pol γ, and PARylated Pol γ should diminish at PARP1 to Pol γ molar ratio ≥1. On the contrary, Pol γ PARylation did occur at the 1:1 molar ratio and was strengthened with increasing concentrations of PARP1 (Fig. 2E).

As PAR chains elongate, the negatively charged polymers cause PARylated proteins to dissociate from DNA. Because PARylated PARP1 displays significantly lower catalytic activity (*28*), the above results suggest that PARP1 either PARylates Pol γ prior to or simultaneously with auto-PARylation before dissociation from DNA. In either case, PARP1 binding rate to the gapped DNA must be faster than Pol γ gap-filling synthesis to halt its activity. The excess non-PARylated PARP1 correlates well with the decreased activity (Fig. 2G), supporting the notion that PARylated PARP1 dissociates from the DNA whereas the non-PARylated species blocks Pol γ activity.

### LC-MS-MS identification of PARylated amino acids

To pinpoint the location of PARylation to provide structural understanding of PARP1-Pol γ interaction, we performed liquid chromatography with tandem mass spectrometry (LC-MS-MS) following the method established by Zhang et al (*27*). This method utilizes hydroxylamine to digest the PAR chains an -NH group on glutamates and aspartates and the peptides and residues containing an additional 15 Da was identified in MS/MS. Duplicated samples were analyzed for duplicated samples containing Pol γ, PARP1 and the 75nt gapped DNA, with or without NAD^+^. The threshold for identification of PARylation sites was set at 2-fold above the corresponding sample without NAD^+^.

The peptides from catalytic subunit Pol γA, accessory subunit Pol γB and PARP1 were recovered at 87%, 75%, and 82%, respectively. PARylation sites on Pol γA and Pol γB as well as PARP1 were all detected. Comparison to the sample without NAD^+^, samples with NAD^+^ displayed increased PARylation sites on Pol γA, Pol γB, and PARP1, which include 12 on PARP1, 16 on Pol γA, and 5 on Pol γB, suggesting the modification sites are correlated with PARylation (Table S3). The detected PARP1 modification sites are a subset of those previously reported from cell lysate (*27, 29*), perhaps due to the difference in detection thresholds or in purified proteins vs more complex cellular environment. Although the method detects PARylation only on glutamate and aspartate residues, the relative PARylation of Pol γ to PARP1 are consistent with that detected by Western Blot (Fig. 1). To visualize the location of the PARylation sites, we composed a model of the Polγ-PARP1-gap DNA complex using crystal structures of Pol γ-DNA (PDB: 4ZTZ) and PARP1-DNA (PDB: 3ODC, 4DQY) by aligning their respective DNA. In the model, Pol γA is in close contact with PARP1 at the 3’-end and 5’-end of the DNA gap and Pol γB is distal to PARP1. When PARylation sites were mapped on the model, most Pol γA PARylation sites are near PARP1. Considering Pol γ does not have canonical PAR binding motif, the results suggest Pol γ PARylation is enhanced by its close vicinity of PARP1 on the gapped DNA (Fig. 2H).

Pol γB is a dimer, the LC-MS/MS revealed sites cannot be unambiguously identified on each monomer. The number of PARylation sites was then estimated using ^32^P-NAD^+^ in the PARylation reaction followed by enzymatic PAR digestion with human poly(ADP-Ribose) glycohydrolase (hPARG) that leaves the terminal ^32^P-ADP-Ribose monomer (MAR) on the modified amino acids. The intensity of Pol γA is twice that of Pol γB (Fig. EV5). Considering that the number of amino acids in Pol γA and dimeric Pol γB are comparable, the results indicate that only one monomer of Pol γB, likely the proximal monomer, was modified.

### PARP1 has no effect on Pol γ during DNA replication

With the discovery that PARP1 regulates Pol γ DNA repair activity, we next tested effect of PARP1 on Pol γ DNA replication. The assay was carried out under conditions identical to the gap-filling synthesis experiments except that the DNA substrate is the primer/template, 25/45nt. In contrast to PARP1’s regulation of Pol γ gap-filling activity, PARP1 displayed little effect on Pol γ during replication (Fig. 3). Increasing NAD^+^ concentration had no impact on Pol γ activity.

**Figure 3.**
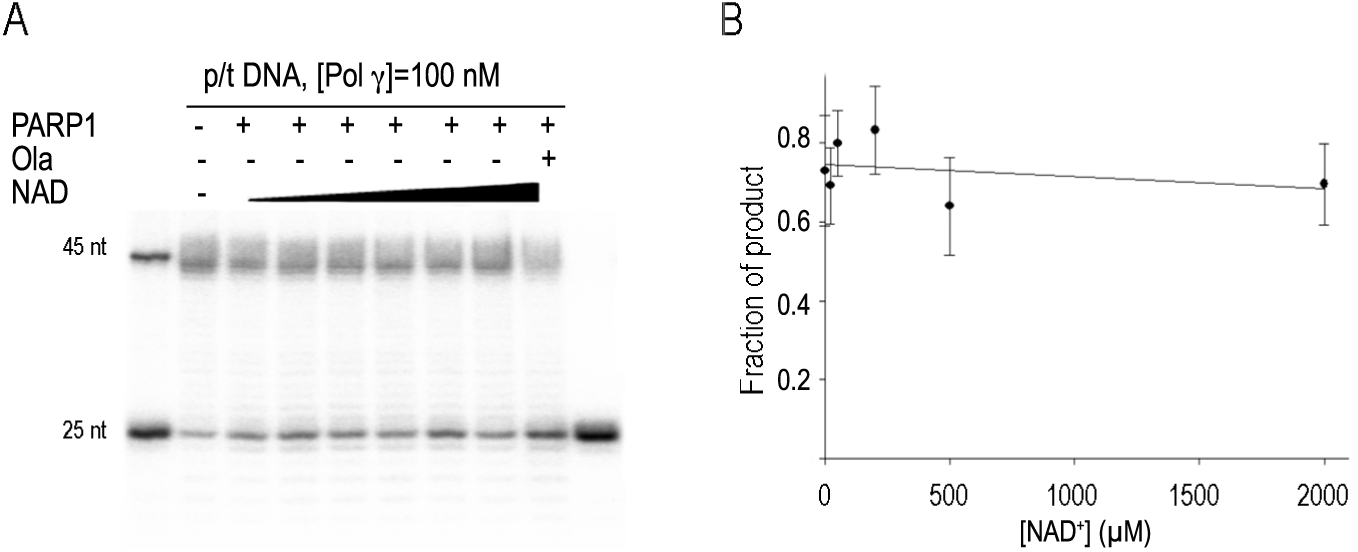
PARP1 effect on Pol γ replication A) Pol γ synthesis on primer/template (25/45nt) DNA in the presence of NAD+, PARP1 and Olaparib. B) quantification.

## DISCUSSION

The closeness of the mitochondrial genome to the OXPHOS electron transfer chain causes more oxidative mtDNA damage by reactive oxygen species (ROS). The levels mtDNA damage in the heart and brain are negatively correlated with longevity in mammals, but such correlation was not observed for nuclear DNA (*30*). MtDNA damage repair has therefore profound impact in ageing process and neurodegeneration diseases.

Despite PARP1’s reported conflicting roles on mtDNA repair (*12*) (*13*) (*14*). PARP1 is consistently found to interact with Pol γ together with other repair enzymes (*12, 15*). As an initial effort to tease out PARP1 functions in mtDNA repair from its activation induced NAD-depletion and consequential cellular dysfunction, we carried out *in vitro* biochemical studies to understand PARP1 and its catalytic activity on Pol γ function in mtDNA repair. Our results show that Pol γ repair activity is strongly regulated by PARP1 in an NAD^+^-dependent manner. Under physiological concentrations of NAD^+^, PARP1 facilitates mtDNA repair, whereas under low NAD^+^ or excessive amount of PARP1, PARP1 inhibits Pol γ repair activity, likely due to persistently binding to the broken DNA ends thus hinders DNA repair. Our findings reconcile seemingly opposing effects of PARP1 on mtDNA repair observed in cellular studies. Depending on the NAD^+^ and PARP1 levels of these *in vivo* assays, PARP1 can be both inhibitory and stimulatory to mtDNA BER.

### Relationship of metabolism and mtDNA repair

We show here that NAD^+^ is a signaling molecule for PARP1 regulation of Pol γ activity. Besides being a substrate for PARylation, NAD^+^ is also a cofactor for glucose and fatty acids metabolic reaction, where foodstuffs catabolic reactions are coupled with NAD^+^ reduction to NADH. Conversion of NADH back to NAD^+^ to ensure continuity of metabolic reactions is accomplished though OXPHOS, where the protons removed from NADH drives the chemical reaction of ATP synthesis. Cellular NAD^+^ level is compartmentalized: mitochondria house the largest NAD^+^ pool at ∼250 µM, and nuclear level is ∼100 µM (*31*) (*32*) (*33*), suggesting larger range of regulation in mitochondria. Under physiological conditions, PARP1 activity is tightly regulated. PARylation is transient in the cell, as PAR chains are rapidly degraded by PARG or other hydrolases (*34*). PARP1 is degraded by caspases, and PARP1 transport into mitochondria are tightly regulated by Mitofilin and other transporters to maintain homeostasis (*35*) (*36*). Overactivation of PARP1 under pathological conditions will inhibit mtDNA damage repair, which, in turn, exacerbates OXPHOS dysfunction and reduces ATP production. Furthermore, PARP1 directly inhibit hexokinase 1 thereby glycolysis (*37*) (*38*), which can further reduce ATP level. Prolonged ATP depletion inevitably leads to necrotic cell death. MtDNA maintenance is intimately linked to cellular NAD^+^ level, metabolism and PARP1 activity.

### PARP1 regulates Pol γ DNA replication and repair differently

Unlike the nuclear DNA polymerases with distinguished roles in DNA replication or repair, Pol γ carries out DNA synthesis for both processes in mitochondria. Our studies revealed PARP1 can distinctively regulates Pol γ activities. While Pol γ function in DNA replication is likely insensitive to PARP1, Pol γ DNA repair function is tightly regulated by PARP1 and proceeds only in the presence of sufficient NAD^+^ and competent PARylation. Our results suggest that while both mtDNA replication and damage repair take place under physiological conditions, only replication continues under conditions where PARP1 are overexpressed and metabolism is low, such as prolonged oxidative stress, ageing process or degenerative diseases. Impediment of mtDNA repair and low metabolism will increase mtDNA mutations and reduced ATP production, which could lead to mitochondria-dependent apoptosis (*39*).

### A model for PARP1-Pol γ interplay in mtDNA repair

Increasing evidence indicates that repair enzymes form a complex, referred to as the repairosome. The DNA repair intermediates are proposed to be handed-off among the enzymes until repair is completed. PARP1 serves as a scaffolding protein for repairosome formation. In line with this theory, we summarize our findings in the following model.

Upon lesion removal and DNA end cleaning, PARP1 binds the 5’-end of the gap and recruits Pol γ to the 3’-end of the gap. Because auto-PARylated drastically reduces PARP1 catalytic activity (*28*), in the absence of NAD+, PARP binding to the 5’-end prevents Pol γ from gap-filling synthesis. In the presence of PARP1, PARP1 catalyzes Pol γ trans-PARylation first or simultaneously with auto-PARylation. The modified PARP1 dissociates from the DNA, thereby removing the physical hindrance to Pol γ, which proceeds with gap-filling synthesis and BER continues (Fig.4).

**Figure 4.**
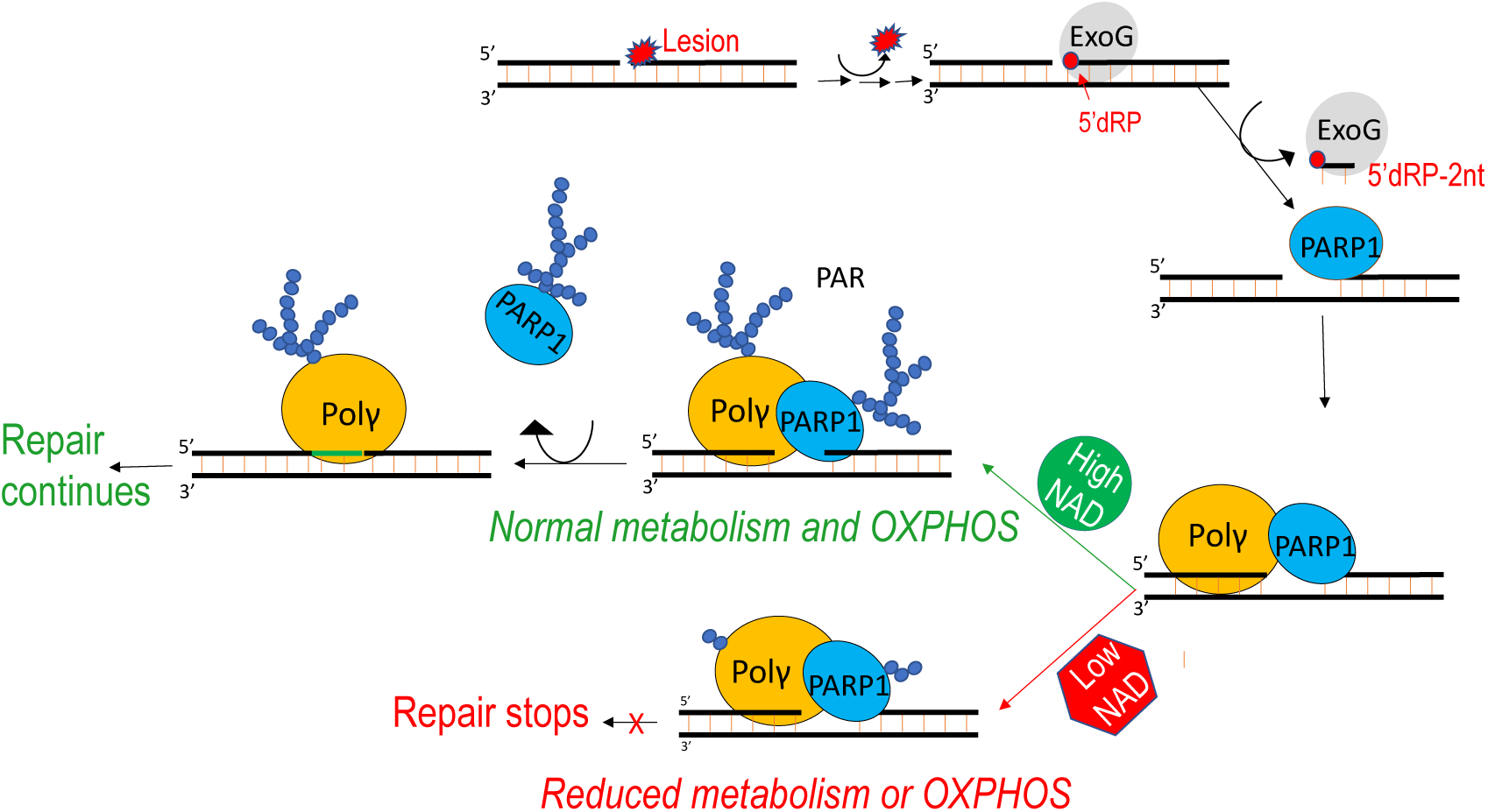
A proposed model for coordinated activity of PARP1 and Pol γ in mitochondrial BER and the association of mtDNA repair and metabolism.

Besides being therapeutics to fight breast and ovarian cancer in DSBR-deficient patients, PARP1 inhibitors are also used in cardiac diseases and infection to suppress overexpressed PARP1. PARP overactivation plays a pivotal role in transformation pathological cardiac hypertrophy to heart failure (*40*). PARP1 inhibitors improve symptoms, nevertheless, their prolonged usage could increase mutation rates (*40*). Our study suggests that inhibition of PARP1 catalytic activity as well as excessive PARP1 halt mtDNA repair and will exacerbate mtDNA damage, explaining at molecular level that prolonged usage of PARP1 inhibitors could stress mitochondria in cardiomyocytes and induce dysfunction. If a drug can be designed to prevent PARP1 binding to DNA rather than PARylation, it may achieve higher efficacy with lower drug toxicity. Additionally, perhaps jointly inhibiting Mitofilin and PARP1 could overcome certain PARP1 side effects in treatment of cardiovascular disorders. As mitochondria-specific PARP1 inhibitors have been synthesized (*41*), they can be used to further tease out PARP1’s function in mitochondria and nucleus.

## MATERIALS AND METHODS

### Preparation of oligonucleotide substrates

Synthetic DNA oligonucleotides were purchased from Integrated DNA Technologies (Coralville, Iowa) or Midland Certified Reagent Company (Midland, Texas). Oligonucleotide sequences are listed in Table S1.

All oligonucleotides were annealed in buffer containing 20 mM Tris (pH 8.0), 100 mM NaCl, 10% glycerol (v/v), 1 mM EDTA by heating to 95°C followed by slow cooling to room temperature. The primer/template P25/T45 was formed with 25nt primer annealed to the 45nt template at 1/1.1 molar ratio, and the 3nt gap substrate T45/P25-B17 was formed by annealing T45 to P25 and a 17nt DNA (B17) that anneals to the 5’end of T45 at 1/1.1/1.2 molar ratio. Self-complementary 75-nt dumbbell with a 3nt gap (d3ntgap) and the 8nt self-annealing DNA were annealed at 10 µM stock concentration (Fig. 1&2)

### Protein purification

Pol γA and Pol γB purification was carried out following the previously published protocol (*1*). Briefly, PolγBΔI4 variant was expressed in *E. coli* BL21-RIL and purified using Ni-NTA agarose (Qiagen) and Mono S affinity chromatography. Pol γA was expressed in sf9 cells and purified on TALON (Clontech) and Superdex200 size exclusion columns. Purified Pol ⍰ A was mixed with Pol γB at a 1:2 molar ratio and applied to the Superdex200 gel filtration column. Peak fractions corresponding to the Pol γAB holoenzyme heterotrimer were pooled and concentrated. Purity was judged to be ∼98% using SDS-PAGE. PARP-1 was expressed in *E. coli* BL21 (DE3) and purified per a published protocol (*2*). Following cell lysis, PARP-1 was purified by sequential application to Ni-NTA agarose (Qiagen), HiTrap Heparin HP column (GE Healthcare), and Superdex200 chromatography columns. Human Poly(ADP-ribose) glycohydrolase catalytic domain (deletion of N-terminal 455 amino acids) (hPARG-ΔN455) was purified according to (*3*). After expression in *E. coli* BL21 (DE3), hPARG-ΔN455 was purified by sequential application to Ni-NTA agarose (Qiagen) and Superdex200 chromatography columns.

### Isothermal Titration Calorimetry

PARP-1 and Pol γ were dialyzed against Buffer RX (25 mM HEPES, pH 7.5, 140 mM KCl, 5 mM MgCl_2_, 5% Glycerol, 1 mM β-mercaptoethanol (BME)) at 4°C overnight. Following dialysis, and and concentrations were determined using their respective extinction coefficients. Titrations were carried out in triplicate (PARP-1 and d3ntgap) or duplicate (PARP-1, Pol γ, and d3ntgap) using a Malvern MicroCal PEAQ-ITC at 25°C with 19 injections (0.4 µL for the 1^st^ injection, followed by 18 injections of 2 µL) of 50 µM d3ntgap titrated into 5 µM protein. For titrations involving both proteins, both were 5 µM. Control titrations were performed by titrating 50 µM d3ntgap into buffer. Thermodynamic binding parameters were determined using NITPIC and SEDPHAT as described in (*4*). Figures for publication were produced using GUSSI (*5*).

### Super-shift Electrophoretic Mobility Shift Assay (EMSA)

Pol γ (0.5 µM) was incubated with increasing concentrations of PARP-1 (0.05 µM, 0.1 µM, 0.2 µM, 0.4 µM, 0.6 µM, 0.8 µM, 1 µM, 1.2 µM, 1.5 µM, and 2 µM) in Buffer RX on ice for 10 minutes. Gapped DNA substrate (d3ntgap, 0.75 µM) was added, and the protein-DNA mixtures were incubated at room temperature for 10 minutes. Samples were loaded onto a gradient 4-20% native PAGE gel (Bio-Rad), electrophoresed at 180V for ∼1hr in 0°C native running buffer (25 mM Tris, 192 mM glycine). Gels were stained in 1X SyBr Gold (Invitrogen) (diluted in native running buffer) and scanned using the Cy2 setting on a Typhoon Amersham Biomolecular Imager. Densitometry was performed with IQTL. The fraction of complex formed was calculated as:

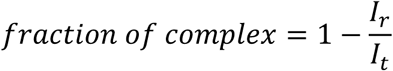

where *I*_*r*_ denotes the intensity of the remainder Po⍰l -DNA band, and *I*_*t*_ denotes the total combined intensities of the Pol γ-DNA band without no PARP1. Due to the diffuse nature of the super-shifted species, Pol γ-PARP1-DNA complex formation is more accurately measured by the disappearance of the Pol γ-DNA binary complex. The fraction of complex formed was plotted as a function of PARP-1 concentration to obtain an apparent dissociation constant Kd.

### Trans-PARylation Assay

Polγ (1 µM) was incubated with d3ntgap (0.75 µM) and increasing concentrations (2 µM, 20 µM, 200 µM, 2000 µM) of NAD^+^ in Buffer RX at 25°C for 5 min prior to addition of PARP1 (1 µM) and incubated at 37°C for 30 minutes. Reactions were quenched with SDS-PAGE loading buffer (60 mM Tris-HCl, pH 6.8, 2% SDS, 10% glycerol, 0.025% bromophenol blue, and 5% BME), and heated at 95°C for 5 min. Products were resolved by electrophoresis on 10% denaturing PAGE and analyzed by Western blot

Gels were incubated in transfer buffer (25 mM Tris base, 192 mM glycine, 10% methanol) at 25°C for 10 min with gentle agitation. Reaction products were transferred to nitrocellulose membranes at 100V for 70 min in transfer buffer; the membranes were then washed with 1X TBST (20 mM Tris, 150 mM NaCl, 0.05% Tween 20 (v/v)) and blocked using 1X TBST supplemented with 5% (w/v) skim milk powder for 1 hr at room temperature. Membranes were then sequentially probed for 1 hr with primary and secondary antibodies (details in Table S2) at room temperature. Between probes, membranes were thoroughly washed with 1X TBST. Products were visualized by chemiluminescence (Pierce^™^ ECL Western Blotting Substrate) using an ImageQuant LAS 4000.

Quantification of PARylated was calculated from the residual of non-PARylated species (I_protein_residual_) normalized by the control intensity within the lane (I_protein _total_)

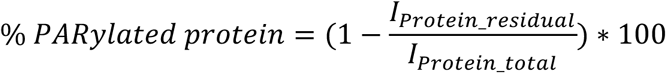

### PARylation sites number determination

PAR chains on Pol Ill or PARP1 were converted to mono(ADP-ribose) (MAR) by carrying out trans-PARylation assays similar to the above assay except that 200 µM ^32^P NAD^+^ (1.6 µCi,) was used. PARylation was stopped by addition of Olaparib (20 µM) followed by the addition of hPARG-ΔN455 (5 µM). Reaction mixtures were incubated at 37°C for 30 min, then quenched with an equal volume of SDS-PAGE loading buffer and resolved on 10% denaturing PAGE. After autoradiography, products were visualized using an Amersham Typhoon Biomolecular imager (GE) and quantified using IQTL.

### DNA synthesis activity assays

DNA replication and gap-filling synthesis were carried out under identical conditions except the DNA substrate. A primer/template duplex (5’-^32^P-25nt/45nt) was used for replication and a 3nt gapped DNA (T45/5’-^32^P25-B17) for gap-filling repair activity. 100 nM Pol 1 and 200 nM PARP-1 (where indicated) were mixed with 100 nM DNA substrate in Binding Buffer (25 mM HEPES, pH 7.5, 140 mM KCl, 5% glycerol, 0.5 µg/mL BSA, 1 mM BME). Incubation was at 0°C for 10 min and then at room temperature for 5 min. Where indicated, Olaparib (10 µM) was added during the room temperature incubation, prior to reaction initiation. PARylation and DNA synthesis were initiated by addition of NAD+, 10 mM MgCl_2_ and 50 µM dNTP. Reaction mixtures were incubated at 37°C for 5 minutes, quenched with stop buffer (80% formamide, 50mM EDTA, 0.1% SDS, 5% glycerol, 0.02% bromophenol blue), heated at 95°C for 5 min, separated on 20% PAGE/7M Urea gels in 1X TBE at 15W for ∼1.5 hrs and autoradiographed. The substrate and product were visualized on an Amersham Typhoon Biomolecular imager (GE) and quantified using ImageQuant. Fraction of product was calculated using the intensities of DNA substrate (I_substrate_) and product (I_product_)

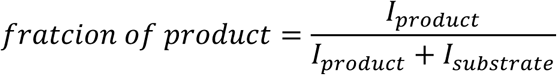

### Liquid Chromatography tandem mass spectrometry (LC-MS/MS)

The trans-PARylated proteins were denatured in 1% SDS at room temperature. To each 22 µL denatured sample, an equal volume of 10% SDS in 100 mM triethylammonium bicarbonate (TEAB) buffer (pH 8.0) was added, followed by 1 µL of 0.25M Bond-Breaker TCEP solution (ThermoFisher). The mixture was incubated at 56°C for 30 min. After samples cooled to room temperature, 0.5M Iodoacetamide (BioUltra) was added and incubated at room temperature for 30 minutes in the dark. 2.7 µL of 12% phosphoric acid was added in the dark, followed by 165 µL of 90% methanol in 100 mM TEAB buffer. The solution was applied to a micro S-Trap column and centrifuged for 1 min at 1,200 cpm. The S-Trap was then washed sequentially with 100 mM TEAB buffer, 90% methanol in 100 mM HEPES (pH 8.5), and 150 mM NaCl in 100 mM HEPES (pH 8.5). 0.5M NH_2_OH was applied to the S-Trap and incubated at room temperature overnight. The S-trap was washed twice with TEAB buffer before adding 25 µL of 20 ng/µL Trypsin (in 50 mM TEAB) and incubating at 47°C for 2 hours. Tryptic peptides were eluted from the S-Trap with 40 µL of 50 mM TEAB, 40 µL of 0.2% formic acid (FA), 30 µL of 50% acetonitrile (ACN) with 0.2% FA, and 30 µL of 80% ACN with 0.1% FA. The eluted tryptic peptides were dried in a speed vac and reconstituted with 100 µL of 2% ACN and 0.1% FA for mass spectrometry.

Nano-LC/MS/MS was performed on a Thermo Scientific Orbitrap Fusion system coupled with a Dionex Ultimat 3000 nano HPLC and auto sampler with 40 well standard trays. The sample was injected onto a trap column (300 µm i.d. x 5 mm, C18 Pep Map 100, Thermo) followed by a C18 reverse-phase nano LC column (Acclaim PepMap 100 75 µm x 25cm, Thermo), both of which had previously been heated to 50°C in the chamber. The flow rate was set to 400 nL/min with a 60-min gradient, where mobile phases were A (99.9% water, 0.1% FA) and B (99.9% ACN, 0.1% FA). As peptides eluted from the column, they were sprayed through a charged metal emitter tip into the mass spectrometer. Parameters included the following: tip voltage + 2.2 kV, FTMS mode was set for MS acquisition of precursor ions (resolution 120,000); ITMS mode was set for subsequent MS/MS via HCD on top speed every 3 sec.

Proteome Discoverer 1.4 was used for protein identification and peak area detection. The UniProt human database was used to analyze raw data. The digestion enzyme was set to Trypsin, Carbamidomethyl Cysteine was as a fixed modification, oxidized methionine and PARylated asparte and glutamate were set as dynamic modifications. A maximum of 2 missed cleavages was allowed. The precursor mass tolerance was set to 10 ppm, the MS/MS fragment mass tolerance was 0.2 Da, and peptides with +2, +3, and +4 charges were considered. The resulting peptides are considered as significant when the False Discovery Rate is ≤1%.

## ACKNOWLEDGMENTS

We thank Drs. John Pascal, Ivan Ahel, and Wlodek Bujalowski for clones of human PARP1, hPARGΔ1-455, and Pol β and help with purification, Michal Szymanski for purified hEXOG-ΔN58 and Polβ, Ian Molineux, Joon Park and Dr. Katherine Kaus for discussion and critical reading of the manuscript, Dr. Xuemei Luo for assistance with mass spectrometry. The work is supported by grants from NIH (GM 110591 and AI134970 to YWY).

## EXPANDED VIEW

**Table EV1.**
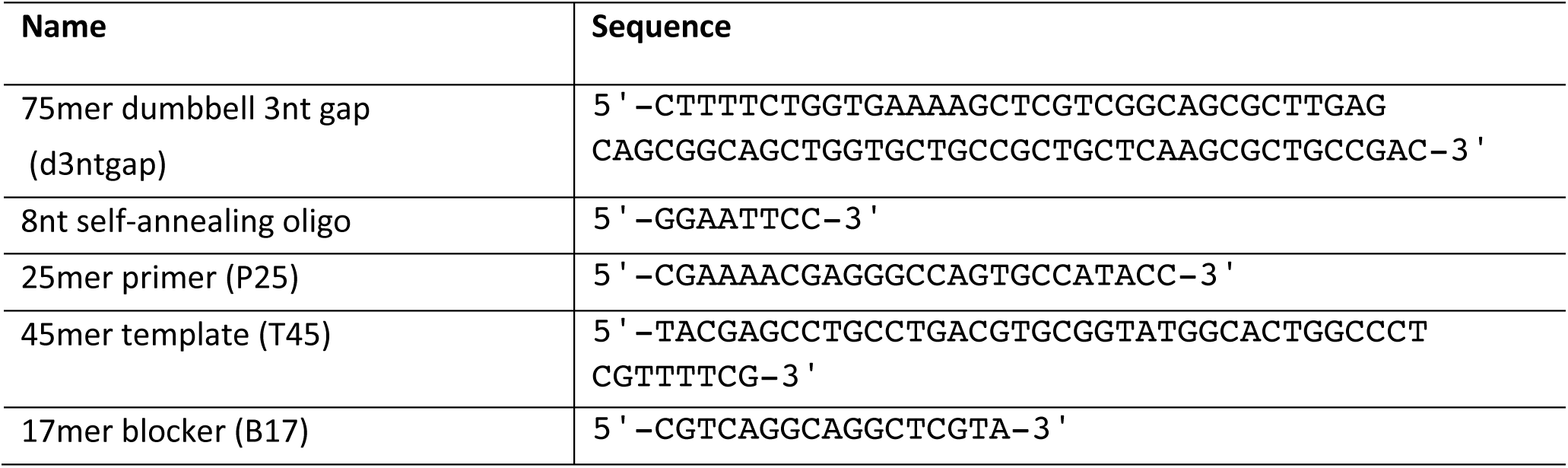
Oligonucleotide sequences

**Table EV2.** Antibodies, sources and titers

| Primary Antibody |  |  | Secondary Antibody |  |  |
| --- | --- | --- | --- | --- | --- |
| Antibodies | Manufacture | Dilution | Host, against | Manufacture | Dilution |
| Mouse, anti-PARP1 | ThermoFisher | 1:2,000 | Goat, anti-mouse | Abcam | 1:5,000 |
| Rabbit, anti-PolyA | ThermoFisher | 1:2,000 | Goat, anti-rabbit | ThermoFisher | 1:5,000 |
| Mouse, anti-PAR | Enzo | 1:2,000 | Goat, anti-mouse | Abcam | 1:5,000 |
| Mouse, anti-His | GE healthcare | 1:3,000 | Goat, anti-mouse | Abcam | 1:5,000 |

**Table EV3.**
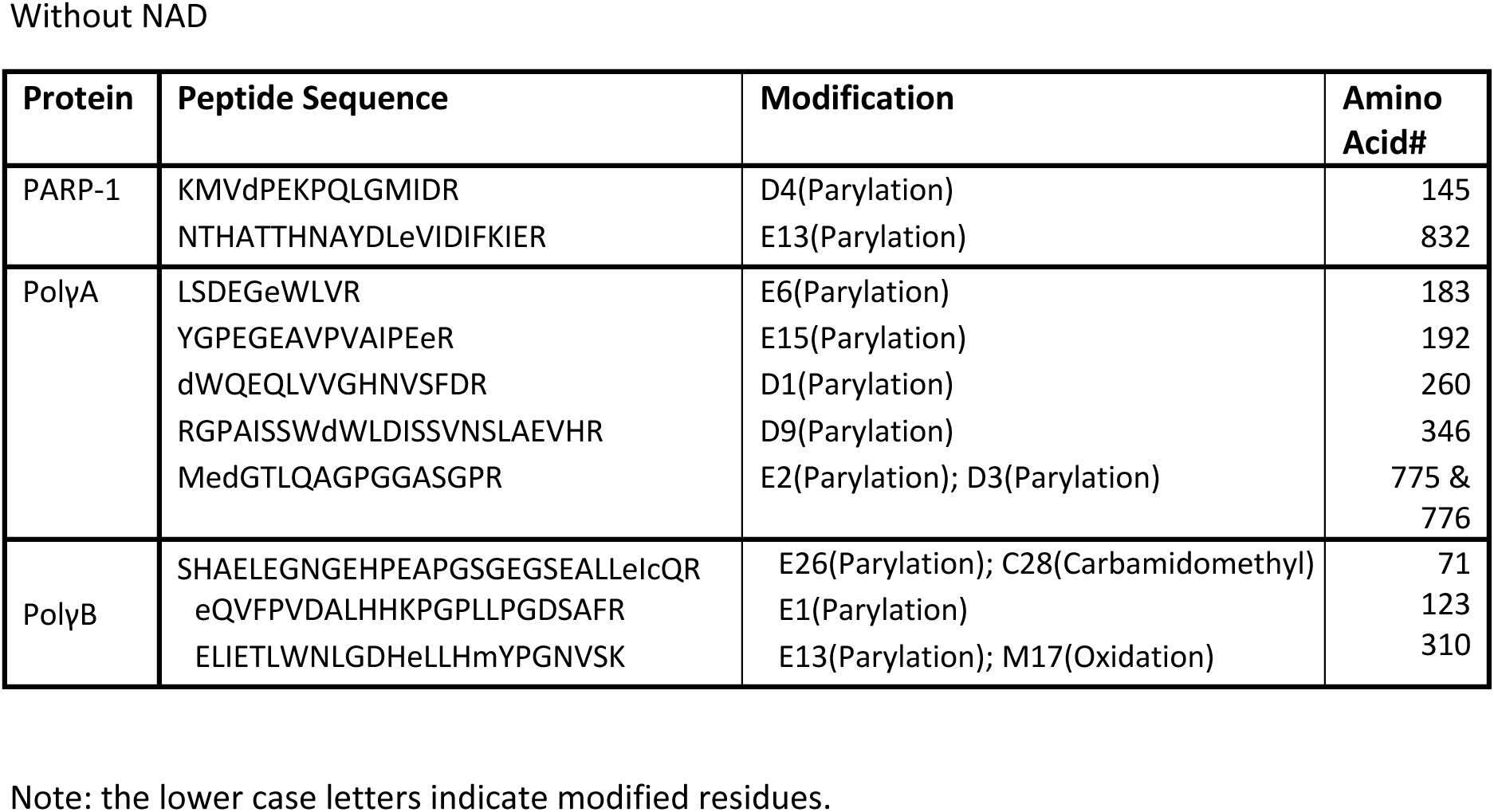
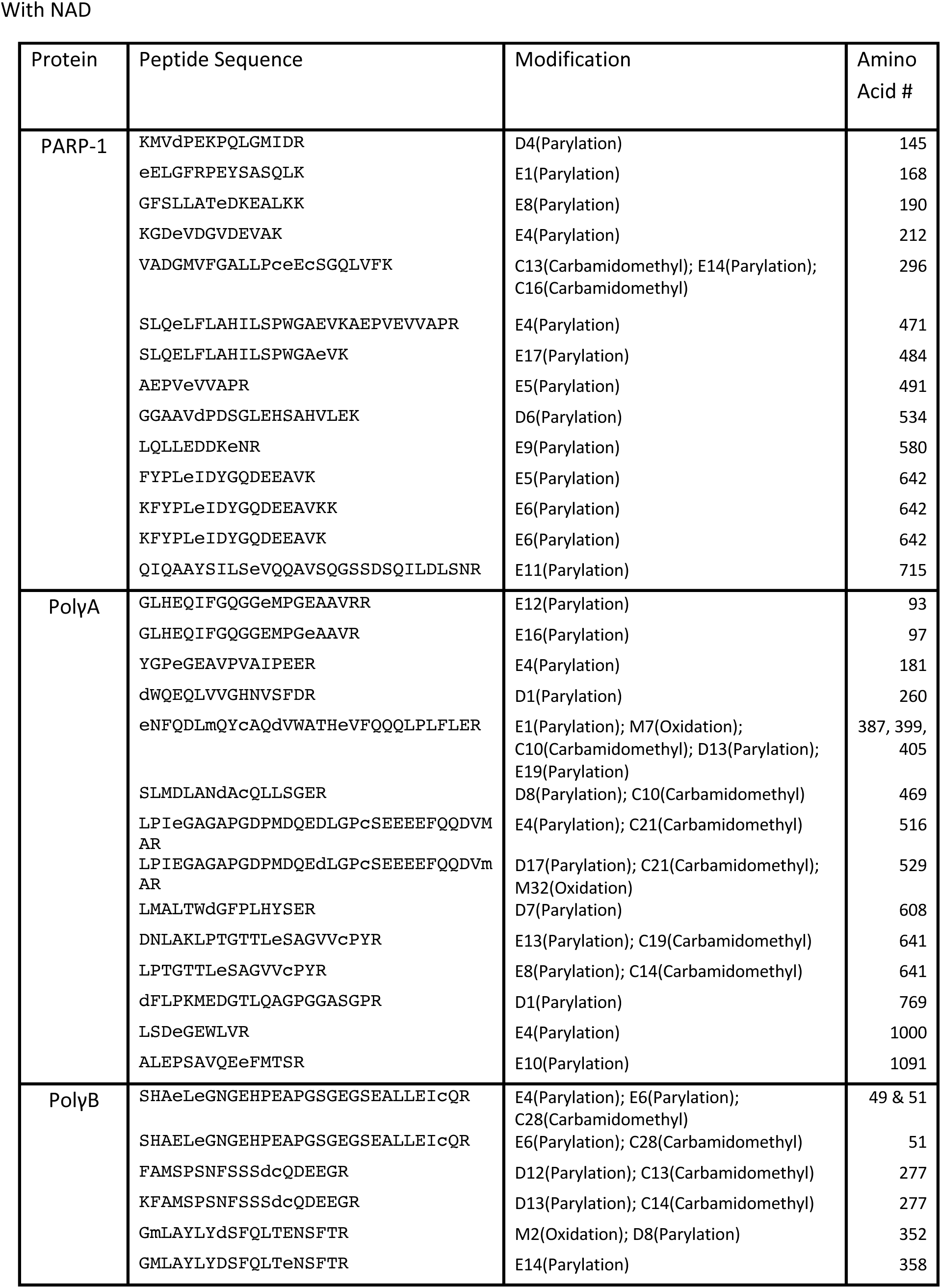
PARylation sites detected by mass spectrometry

## SUPPLEMENT FIGURES

**Figure EV1.**
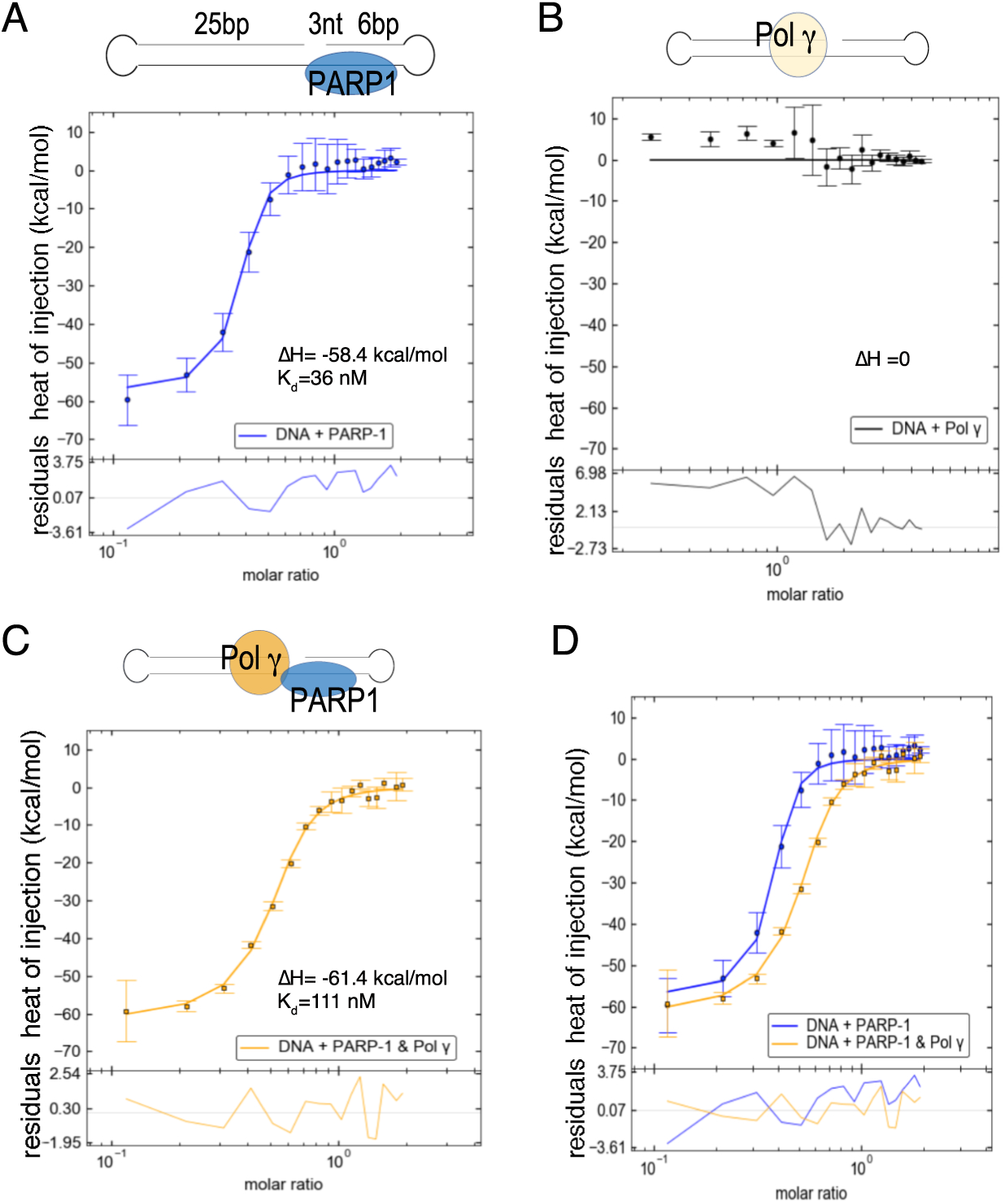
Pol γ-PARP1 interaction. A-C) ITC measurement for gap-DNA binding for PARP1, Pol γ alone and PARP1-Pol γ. D) Superposition of PARP1-DNA and PARP1-Pol γ-DNA complexes showing a consistent enthalpy (ΔH) difference. All measurements were averaged from three independent experiments.

**Figure EV2.**
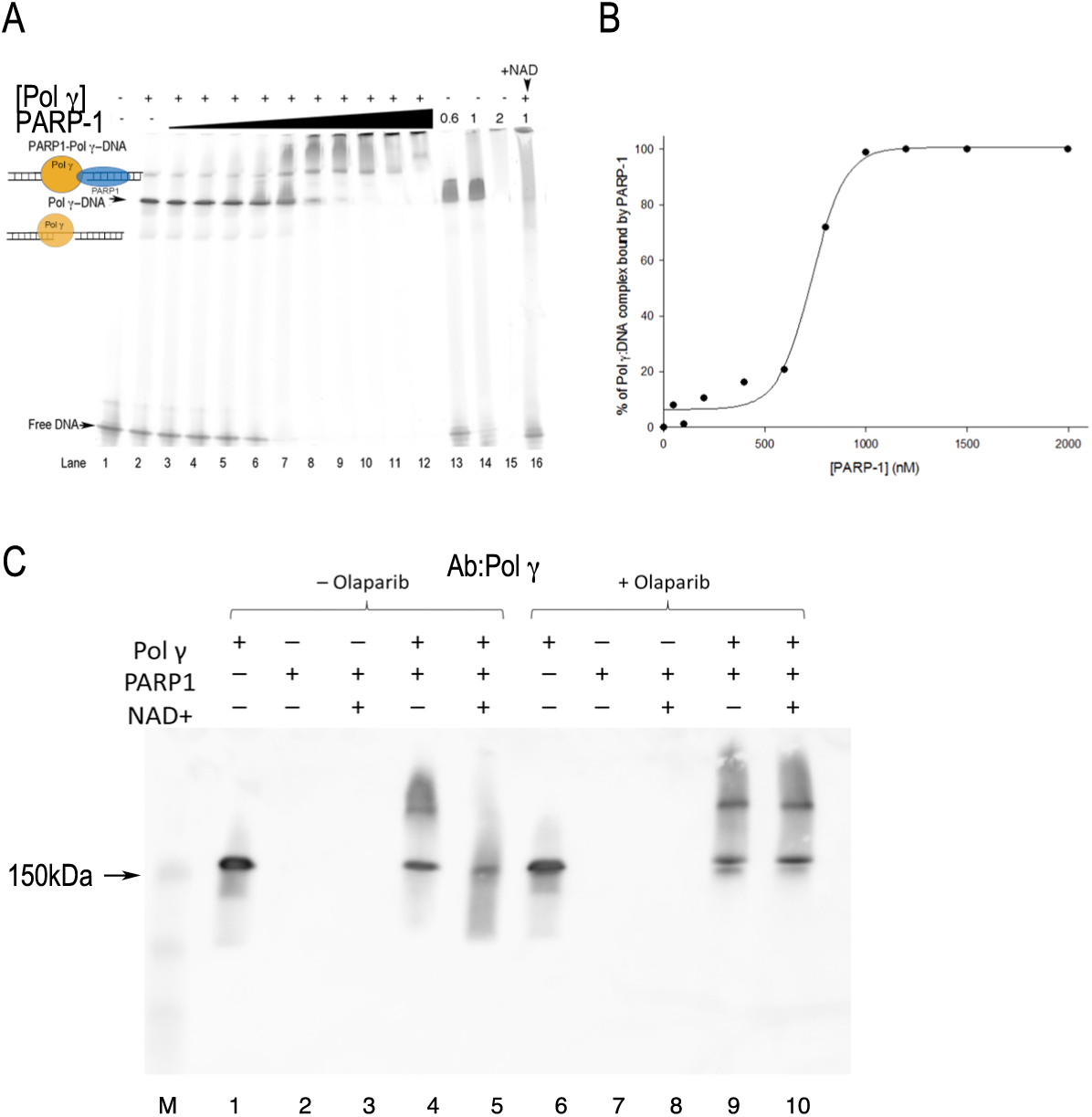
EMSA supershift of Pol γ-DNA-PARP1 complex formation A) Titrating toPol γ-DNA complex with PARP1 at 50, 100, 200, 400, 600, 800, 1000, 1200, 1500, and 2000 nM concentrations. B) quantification of A), where the estimated K_d_ for PARP1-Pol γ-DNA complex is ∼600 nM. C) Western Blot of EMSA with antibody against Pol γ. All samples contain 0.75 µM 3nt-gapped DNA (d3ntgap) with (Lane 1-5) or without Olaparib (Lane 6-10), 0.5 µM Pol γ, 1 µM PARP1 and 2 mM NAD^+^. Lane M is a size marker.

**Figure EV3.**
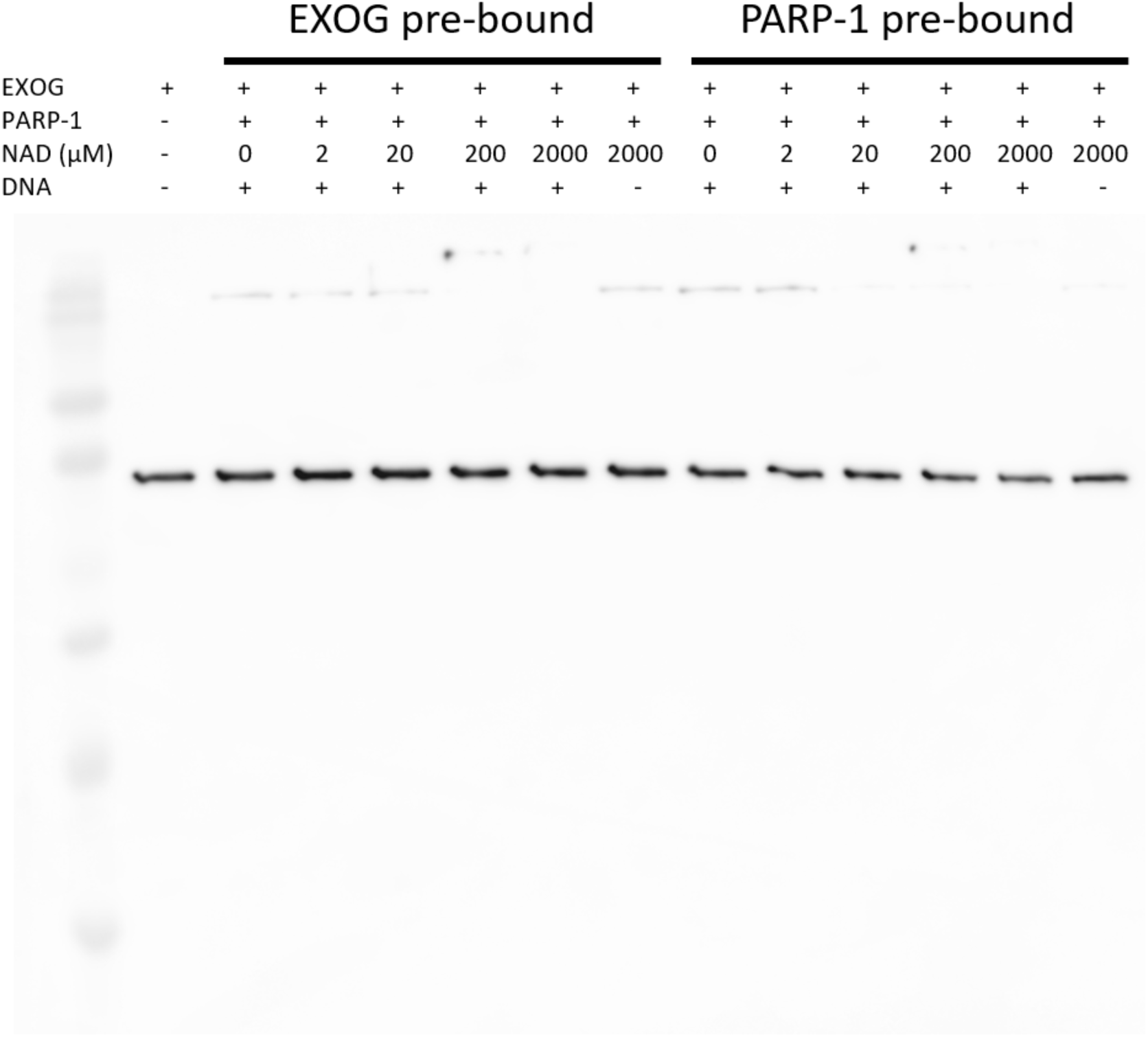
PARylation of ExoG on gapped-DNA Western blot of PARP1 trans-PARylation of catalytic inactive His-EXOG H140A on a gapped DNA at increasing concentrations of NAD+ that was probed with anti-His antibodies. The orders of EXOG and PARP1 addition to the DNA were tested.

**Figure EV4.**
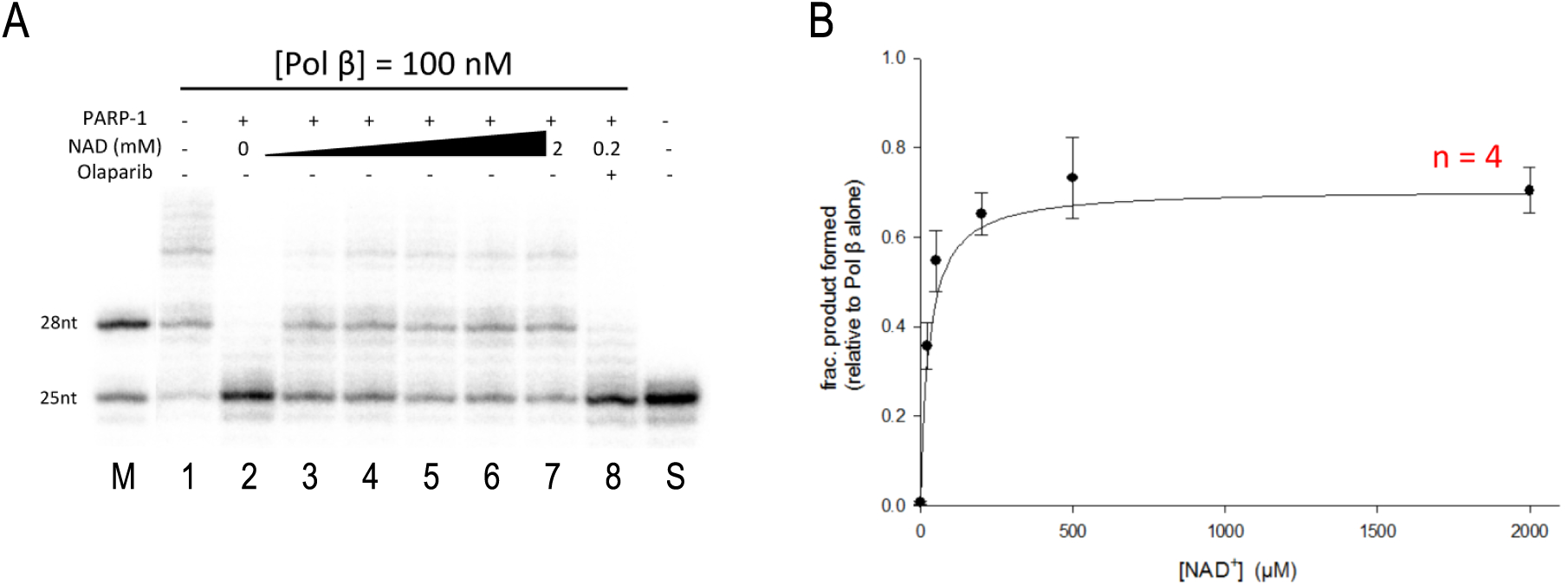
Effect of PARP on Pol β gap-filling activity. A) Pol β activity was analyzed in the absence 0f PARP1 (Lane 1), in the presence of PARP (Lane 2) and at increasing concentrations of NAD (Lanes 3-7), as well as with PARP1 inhibitor, Olaparib (Lane 8). Lane M and S are marker and substrate only, respectively. B) quantification of Pol b product as a function of NAD (plotted the mean of four experiments with standard deviation)

**Figure EV5.**
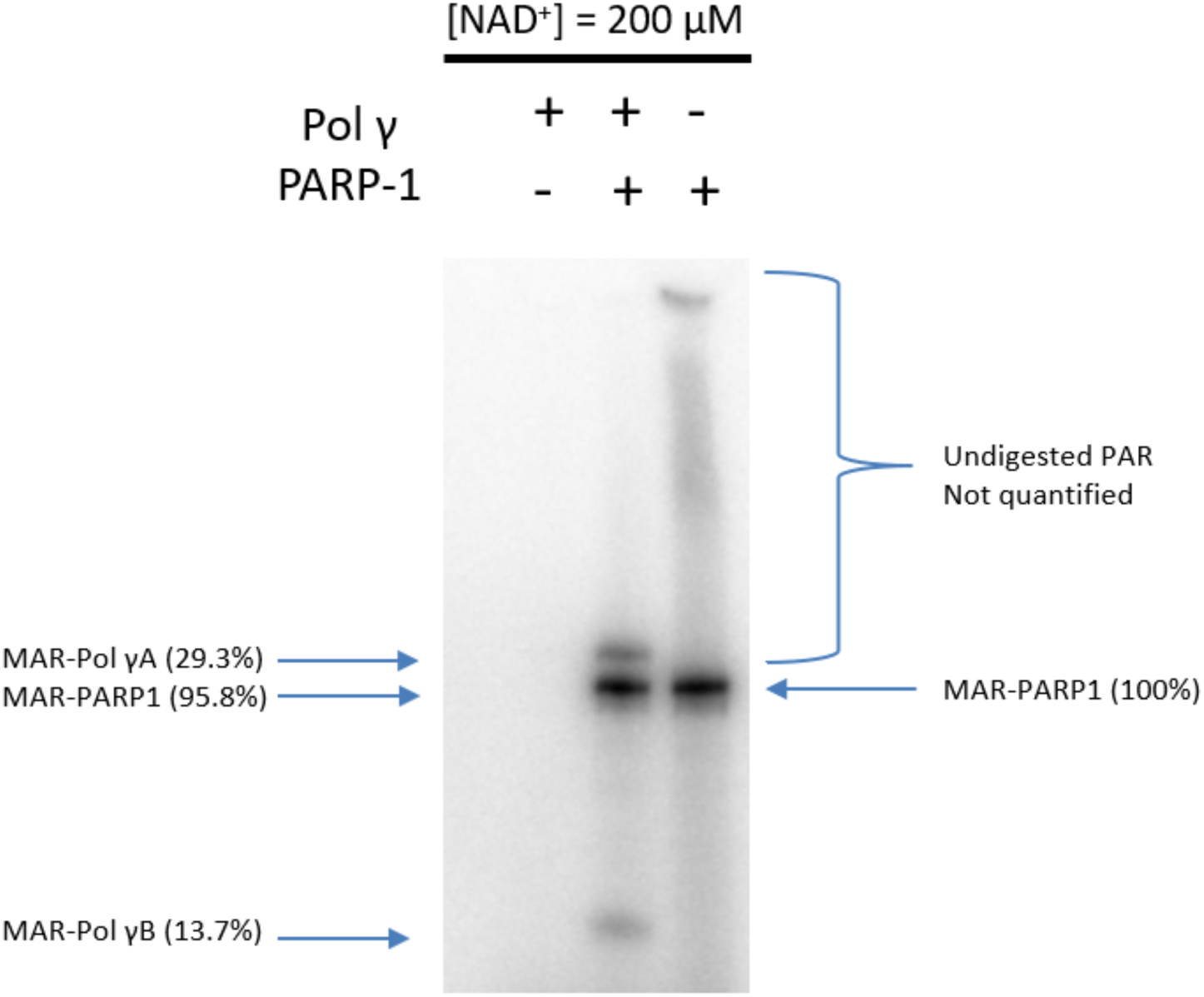
Determination of auto- and trans-PARylation sites numbers. Mono (^32^P-ADP-ribose) (MAR) modified Pol γA, Pol γB and PARP1 quantified and normalized to the density of the MAR-PARP1 band in lane 3 (PARP1 alone).

## REFERENCES

1. W. M. Brown, M. George, Jr., A. C. Wilson, Rapid evolution of animal mitochondrial DNA. Proc Natl Acad Sci U S A 76, 1967–1971 (1979).

2. W. M. Brown, E. M. Prager, A. Wang, A. C. Wilson, Mitochondrial DNA sequences of primates: tempo and mode of evolution. J Mol Evol 18, 225–239 (1982).

3. K. H. Almeida, R. W. Sobol, A unified view of base excision repair: lesion-dependent protein complexes regulated by post-translational modification. DNA Repair (Amst) 6, 695–711 (2007).

4. W. C. Copeland, The mitochondrial DNA polymerase in health and disease. Subcell Biochem 50, 211–222 (2010).

5. Q. He, C. K. Shumate, M. A. White, I. J. Molineux, Y. W. Yin, Exonuclease of human DNA polymerase gamma disengages its strand displacement function. Mitochondrion 13, 592–601 (2013).

6. I. A. Cymerman, I. Chung, B. M. Beckmann, J. M. Bujnicki, G. Meiss, EXOG, a novel paralog of Endonuclease G in higher eukaryotes. Nucleic Acids Res 36, 1369–1379 (2008).

7. M. R. Szymanski et al., A domain in human EXOG converts apoptotic endonuclease to DNA-repair exonuclease. Nat Commun 8, 14959 (2017).

8. A. E. Fisher, H. Hochegger, S. Takeda, K. W. Caldecott, Poly(ADP-ribose) polymerase 1 accelerates single-strand break repair in concert with poly(ADP-ribose) glycohydrolase. Mol Cell Biol 27, 5597–5605 (2007).

9. F. Dantzer et al., Base excision repair is impaired in mammalian cells lacking Poly(ADP-ribose) polymerase-1. Biochemistry 39, 7559–7569 (2000).

10. M. Masson et al., XRCC1 is specifically associated with poly(ADP-ribose) polymerase and negatively regulates its activity following DNA damage. Mol Cell Biol 18, 3563–3571 (1998).

11. O. I. Lavrik et al., Photoaffinity labeling of mouse fibroblast enzymes by a base excision repair intermediate. Evidence for the role of poly(ADP-ribose) polymerase-1 in DNA repair. J Biol Chem 276, 25541–25548 (2001).

12. J. J. Wen, Y. W. Yin, N. J. Garg, PARP1 depletion improves mitochondrial and heart dysfunction in Chagas disease: Effects on POLG dependent mtDNA maintenance. PLoS Pathog 15, e1007065 (2018).

13. S. G. Jarrett, M. E. Boulton, Poly(ADP-ribose) polymerase offers protection against oxidative and alkylation damage to the nuclear and mitochondrial genomes of the retinal pigment epithelium. Ophthalmic Res 39, 213–223 (2007).

14. C. Hegedus et al., PARP1 Inhibition Augments UVB-Mediated Mitochondrial Changes-Implications for UV-Induced DNA Repair and Photocarcinogenesis. Cancers (Basel) 12, (2019).

15. B. Szczesny, A. Brunyanszki, G. Olah, S. Mitra, C. Szabo, Opposing roles of mitochondrial and nuclear PARP1 in the regulation of mitochondrial and nuclear DNA integrity: implications for the regulation of mitochondrial function. Nucleic Acids Res 42, 13161–13173 (2014).

16. S. H. Wilson, T. A. Kunkel, Passing the baton in base excision repair. Nat Struct Biol 7, 176–178 (2000).

17. P. Pacher, C. Szabo, Role of poly(ADP-ribose) polymerase 1 (PARP-1) in cardiovascular diseases: the therapeutic potential of PARP inhibitors. Cardiovasc Drug Rev 25, 235–260 (2007).

18. M. R. Szymanski et al., Structural basis for processivity and antiviral drug toxicity in human mitochondrial DNA replicase. EMBO J 34, 1959–1970 (2015).

19. M. F. Langelier, J. L. Planck, S. Roy, J. M. Pascal, Structural basis for DNA damage-dependent poly(ADP-ribosyl)ation by human PARP-1. Science 336, 728–732 (2012).

20. M. F. Langelier, J. L. Planck, S. Roy, J. M. Pascal, Crystal structures of poly(ADP-ribose) polymerase-1 (PARP-1) zinc fingers bound to DNA: structural and functional insights into DNA-dependent PARP-1 activity. J Biol Chem 286, 10690–10701 (2011).

21. M. F. Langelier, D. D. Ruhl, J. L. Planck, W. L. Kraus, J. M. Pascal, The Zn3 domain of human poly(ADP-ribose) polymerase-1 (PARP-1) functions in both DNA-dependent poly(ADP-ribose) synthesis activity and chromatin compaction. J Biol Chem 285, 18877–18887 (2010).

22. J. M. Fischer et al., Poly(ADP-ribose)-mediated interplay of XPA and PARP1 leads to reciprocal regulation of protein function. FEBS J 281, 3625–3641 (2014).

23. P. Loetscher, R. Alvarez-Gonzalez, F. R. Althaus, Poly(ADP-ribose) may signal changing metabolic conditions to the chromatin of mammalian cells. Proc Natl Acad Sci U S A 84, 1286–1289 (1987).

24. Q. Zhang, D. W. Piston, R. H. Goodman, Regulation of corepressor function by nuclear NADH. Science 295, 1895–1897 (2002).

25. M. V. Sukhanova, S. N. Khodyreva, O. I. Lavrik, Poly(ADP-ribose) polymerase-1 inhibits strand-displacement synthesis of DNA catalyzed by DNA polymerase beta. Biochemistry (Mosc) 69, 558–568 (2004).

26. D. D’Amours, S. Desnoyers, I. D’Silva, G. G. Poirier, Poly(ADP-ribosyl)ation reactions in the regulation of nuclear functions. Biochem J 342 (Pt 2), 249–268 (1999).

27. Y. Zhang, J. Wang, M. Ding, Y. Yu, Site-specific characterization of the Asp- and Glu-ADP-ribosylated proteome. Nat Methods 10, 981–984 (2013).

28. P. Zahradka, K. Ebisuzaki, A shuttle mechanism for DNA-protein interactions. The regulation of poly(ADP-ribose) polymerase. Eur J Biochem 127, 579–585 (1982).

29. K. Kalesh et al., An Integrated Chemical Proteomics Approach for Quantitative Profiling of Intracellular ADP-Ribosylation. Sci Rep 9, 6655 (2019).

30. G. Barja, A. Herrero, Oxidative damage to mitochondrial DNA is inversely related to maximum life span in the heart and brain of mammals. FASEB J 14, 312–318 (2000).

31. C. C. Alano et al., Differences among cell types in NAD(+) compartmentalization: a comparison of neurons, astrocytes, and cardiac myocytes. J Neurosci Res 85, 3378–3385 (2007).

32. X. A. Cambronne et al., Biosensor reveals multiple sources for mitochondrial NAD(+). Science 352, 1474–1477 (2016).

33. C. Canto, K. J. Menzies, J. Auwerx, NAD(+) Metabolism and the Control of Energy Homeostasis: A Balancing Act between Mitochondria and the Nucleus. Cell Metab 22, 31–53 (2015).

34. J. M. Pascal, T. Ellenberger, The rise and fall of poly(ADP-ribose): An enzymatic perspective. DNA Repair (Amst) 32, 10–16 (2015).

35. G. V. Chaitanya, A. J. Steven, P. P. Babu, PARP-1 cleavage fragments: signatures of cell-death proteases in neurodegeneration. Cell Commun Signal 8, 31 (2010).

36. M. N. Rossi et al., Mitochondrial localization of PARP-1 requires interaction with mitofilin and is involved in the maintenance of mitochondrial DNA integrity. J Biol Chem 284, 31616–31624 (2009).

37. S. A. Andrabi et al., Poly(ADP-ribose) polymerase-dependent energy depletion occurs through inhibition of glycolysis. Proc Natl Acad Sci U S A 111, 10209–10214 (2014).

38. E. Fouquerel et al., ARTD1/PARP1 negatively regulates glycolysis by inhibiting hexokinase 1 independent of NAD+ depletion. Cell Rep 8, 1819–1831 (2014).

39. B. Van Houten, S. E. Hunter, J. N. Meyer, Mitochondrial DNA damage induced autophagy, cell death, and disease. Front Biosci (Landmark Ed) 21, 42–54 (2016).

40. R. J. Henning, M. Bourgeois, R. D. Harbison, Poly(ADP-ribose) Polymerase (PARP) and PARP Inhibitors: Mechanisms of Action and Role in Cardiovascular Disorders. Cardiovasc Toxicol 18, 493–506 (2018).

41. T. Krainz et al., Synthesis and Evaluation of a Mitochondria-Targeting Poly(ADP-ribose) Polymerase-1 Inhibitor. ACS Chem Biol 13, 2868–2879 (2018).

